# Genomic footprints of historical introgression between ancient lineages of wild *Oryza* AA-genome species with widely separated contemporary distributions

**DOI:** 10.64898/2026.01.14.699414

**Authors:** Kanako O. Koyanagi, Yuta Kotoku, Yuji Kishima

## Abstract

Phylogenetic incongruence is increasingly recognized as pervasive, yet the extent to which reticulate evolution occurs between groups separated by substantial geographical distances and deep phylogenetic divergence remains poorly characterized. In the *Oryza* AA-genome group—a model for plant speciation and domestication—the traditional bifurcation model posits that Australian *Oryza meridionalis* and African *Oryza longistaminata* occupy basal branches, distinct from the more recently diversified monophyletic clade comprising Asian and other African lineages, including major cultivars. However, recent evidence from endogenous viral sequences has hinted at unexpected genetic relatedness between African *O. longistaminata* and Asian *Oryza sativa*, which are geographically and phylogenetically distant. Here, we conducted a genome-wide survey across 11 *Oryza* species to systematically identify genomic regions exhibiting phylogenetic incongruence. Widespread phylogenetic discordance was observed, notably involving genomic segments in which *O. longistaminata* showed phylogenetic proximity to Asian species, contradicting their established deep divergence. To distinguish between introgression and incomplete lineage sorting, we performed four-taxon ABBA-BABA tests, which provided statistical support for introgression. Furthermore, divergence time estimates for these incongruent regions were younger than the species divergence times, suggesting historical introgression between the ancestors of lineages that are currently separated by vast geographical distances. Systematic assessments indicated that potential analytical artifacts, such as compositional bias and substitution saturation, were unlikely to explain the observations. These convergent lines of evidence suggest that ancient introgression had occurred between currently geographically separated and evolutionarily divergent *Oryza* lineages, leaving detectable footprints across their modern genomes.

## Introduction

In the era of phylogenomics, phylogenetic incongruence—the inference of conflicting tree topologies—has become increasingly recognized as pervasive, which can be attributed to both biological and analytical factors (Fleming et al. 2023; Steenwyk et al. 2023). Among the biological factors, incomplete lineage sorting (ILS) and gene flow—such as that caused by hybridization or horizontal gene transfer—are considered major contributors. These processes complicate the reconstruction of speciation process but also enrich our understanding of how species diversify.

Rice (*Oryza sativa*), one of the world’s most important staple crops (Khush et al. 2001; Wang and Han 2022), provides a particularly informative system for studying phylogenetic incongruence, as reported previously (Zou et al. 2008; Cranston et al. 2009; Yang et al. 2011). The genus *Oryza* is characterized by rapid diversification (Zou et al. 2008; Zou et al. 2013; Zhang et al. 2014; Lin et al. 2015; Stein et al. 2018), which can lead to ILS, and phylogenetic incongruence between nuclear and organellar genomes has been reported (Duan et al. 2007; Kim et al. 2015; Yin et al. 2015; Fornasiero et al. 2025), suggesting a complex evolutionary history of these species. In addition, introgression has been documented, for example, between Asian cultivars during domestication contributing to agronomically important traits (Izawa 2022; Zhou et al. 2022; Zhou et al. 2025). However, the extent and causes of phylogenetic incongruence especially among the genomes of wild species remain largely unexplored.

The genus *Oryza* comprises 24 species, including two cultivated rice species, *O. sativa* and *O. glaberrima* (Vaughan et al. 2003; Jena 2010). These two species were domesticated independently from wild species: *O. sativa* from *O. rufipogon* and *O. nivara* in Asia, and *O. glaberrima* from *O. barthii* in Africa (Chang 1976; Vaughan et al. 2008), all of which belong to AA genome group (comprising eight species in total). Because genetic divergence within cultivated rice is limited, its wild relatives, particularly AA-genome species serve as a valuable reservoir of genetic diversity for breeding more resilient and productive varieties (Jacquemin et al. 2013; Brozynska et al. 2016), owing to their relatively low hybridization barriers with cultivated species compared with other genome groups (Zhang et al. 2022; Zhou et al. 2022; Zhou et al. 2025). For example, *Xa21* disease resistant gene from *O. longistaminata* of AA-genome species was utilized to develop bacterial blight resistant rice varieties (Ronald et al. 1992; Brar and Khush 1997). Therefore, understanding evolutionary histories of AA-genome species is of importance.

Phylogenetic relationships among *Oryza* AA-genome species have been studied for decades, and recent whole-genome data have revealed a consensus view: Asian and African (except *O. longistaminata*) species are monophyletic, whereas the Australian *O. meridionalis* and the African *O. longistaminata* fall on deep, distant branches of the AA genome tree (Zhang et al. 2014; Stein et al. 2018; Fornasiero et al. 2025). Today, wild *Oryza* AA-genome species are distributed across tropical and subtropical regions: most species are endemic to specific regions including Asia (*O. rufipogon* and *O. nivara*), Australia (*O. rufipogon* and *O. meridionalis*), Africa (*O. barthii* and *O. longistaminata*), and America (*O. glumaepatula*) (Lu et al. 2000; Atwell et al. 2014). Interestingly, a recent analysis of endogenous rice tungro bacilliform virus-like sequences (ERTBVs) in rice genomes uncovered an unexpectedly close relationship between *O. longistaminata* and *O. sativa* ssp*. japonica*, which are geographically and phylogenetically distant (Saito et al. 2023). This finding challenges the simple bifurcation model of the AA-genome species and raises the question of whether such incongruent patterns extend beyond ERTBV loci in the nuclear genome.

Here, we conducted a genome-wide survey to identify genomic regions exhibiting phylogenetic incongruence among *Oryza* AA-genome species, with a particular focus on phylogenetically divergent AA-genome species, *O. meridionalis* and *O. longistaminata*. By systematically analyzing such regions, we identified signatures of possible ancient introgression between deeply diverged AA-genome species. Notably, these signals involved historical introgression between ancient lineages of *O. longistaminata* and Asian species, currently distributed on different continents.

## Materials and Methods

### Genome data acquisition and processing

We obtained a whole-genome multiple alignment of nucleotide sequences for 11 *Oryza* species representing AA, BB, and FF genome groups from Ensembl Plants release 59 in Multiple Alignment Format (MAF) (https://ftp.ebi.ac.uk/ensemblgenomes/pub/plants/release-59/maf/ensembl-compara/multiple_alignments/11_rice.epo_extended/) (Yates et al. 2022).

The genome assemblies of the 11 species included in the alignment are IRGSP-1.0 for *Oryza sativa* Japonica Group (AA-genome species), ASM465v1 for *Oryza sativa* Indica Group (AA), Oryza_nivara_v1.0 for *Oryza nivara* (AA), OR_W1943 for *Oryza rufipogon* (AA), Oryza_glumaepatula_v1.5 for *Oryza glumaepatula* (AA), O.barthii_v1 for *Oryza barthii* (AA), Oryza_glaberrima_V1 for *Oryza glaberrima* (AA), O_longistaminata_v1.0 for *Oryza longistaminata* (AA), Oryza_meridionalis_v1.3 for *Oryza meridionalis* (AA), Oryza_punctata_v1.2 for *Oryza punctata* (BB-genome species), and Oryza_brachyantha.v1.4b for *Oryza brachyantha* (FF-genome species). The alignment is a progressive, whole-genome multiple alignment using a guide tree and synteny information to align orthologous regions across species generated by EPO pipeline (Herrero et al. 2016). In this study, we used only alignments of chromosomes 1 to 12 of the reference genome (IRGSP-1.0), covering a total of 298,077,048 bp comprising 15,460 alignment blocks in total. Multiple alignment files in MAF format were processed using MafFilter v1.3.1 (Dutheil 2020) to retain only alignment blocks containing all 11 *Oryza* species, using strict=yes option. MafFilter was also used to remove any alignment blocks with duplication, using remove_duplicates=yes option. Sites containing gaps or ambiguous nucleotides (Ns) were excluded. The resulting filtered alignment blocks were then used for subsequent analyses.

### Phylogenetic analysis

Phylogenetic analyses were conducted for each alignment block as well as for a concatenated whole-genome alignment. For analysis of each alignment block, we excluded alignment blocks shorter than 1 kb from the analysis to improve the reliability of the results. Phylogenetic trees were inferred with the maximum-likelihood (ML) method using IQ-TREE v2.4.0 (Minh et al. 2020). The best-fit substitution model for each alignment block was selected by ModelFinder (Kalyaanamoorthy et al. 2017), with -m MFP+LM option. Branch support was assessed using ultrafast bootstrap (Hoang et al. 2018) with 1000 replicates (-B 1000).

For assessing violations of phylogenetic assumptions, a symmetry test was performed using --symtest-only option of IQ-TREE for each alignment block. We also performed a chi-squared test for homogeneity of nucleotide composition for all sequences in each alignment block with IQ-TREE.

Divergence times were estimated using two approaches. First, relative divergence times were estimated using RelTime-ML method (Tamura et al. 2012) implemented in MEGA version 11.0.13 (Tamura et al. 2021) under the GTR+G+I model with five discrete gamma rate categories using non-coding nucleotide sequence setting. Second, absolute divergence times were estimated using a Bayesian approach with BEAST2 v2.7.7 (Bouckaert et al. 2019). We employed the namedExtended model set for a substitution model test, the optimized relaxed clock model with a mean clock rate 6.5x10^-9^ substitutions/site/year (Gaut et al. 1996), and the calibrated Yule model for the tree prior. A calibration point of 15 million years ago was assigned to the AA/FF genome divergence (Tang et al. 2010; Stein et al. 2018), using normal priors with 1.0 sigma based on TimeTree information (Kumar et al. 2022). The Markov Chain Monte Carlo (MCMC) chain length was 5 to 40 (until convergence) million generations, with sampling every 5000 generations. Effective sample sizes (ESS) were initially evaluated using the coda package (Plummer et al. 2006) in R version 4.3.3, and then parameters with ESS ≤ 200 were further inspected in Tracer v1.7.2 (Rambaut et al. 2018). All parameters showed ESS > 200 after confirming convergence. The first 10% of the MCMC samples were discarded as burn-in. Median node ages and 95% highest posterior density (HPD) intervals were generated using TreeAnnotator in BEAST2.

### Detecting introgression

To statistically distinguish introgression from ILS, we performed four-taxon (ABBA-BABA) tests using HYBRIDCHECK (Ward and van Oosterhout 2016) based on the concatenated whole-genome alignment of the alignment blocks. The runForTaxonTests function was performed assuming the tree topology (((P1, P2), P3), A). The four taxa used were *O. punctata* (BB genome species) as ‘A’, either *O. meridionalis* or *O. longistaminata* as P3, and all possible combinations of the remaining species for P1 and P2. To assess statistical significance, the parameter numberOfBlocks was set to 100, and then Patterson’s *D* estimate and its corresponding Z score were calculated by jackknifing the blocks of data. *P*-values were adjusted for multiple testing using the Bonferroni correction.

### Assessment of saturation in nucleotide substitution

To assess the degree of saturation in nucleotide substitution, C (Convergence) value (Struck et al. 2008) was calculated with BaCoCa (Kück and Struck 2014) over all taxa for each alignment block. This value uses the convergence behavior of transition-transversion ratios against the uncorrected genetic p distances. A smaller C value indicates higher degree of convergence and saturation in the alignment (Fleming et al. 2023), although C value does not provide an intrinsic threshold (Struck et al. 2008).

### Validation with other genome assemblies

To assess our findings, we performed sequence similarity searches against several other publicly available genome assemblies. For *O. meridionalis*, haplotype-resolved assembly was downloaded from NCBI Genome database (ASM4749620v1) (Abdullah et al. 2025). Genome assemblies of *O. longistaminata* were downloaded from https://ftp2.cngb.org/pub/CNSA/data5/CNP0005176/CNS0677859/CNA0105971/ for T2T assembly (Guang et al. 2025), from https://pr-nxy.ynu.edu.cn/OLongiDB/download.php for haplotype-resolved assembly (Lian et al. 2024), and from http://www.olinfres.nig.ac.jp/ for Oryza_longistaminata_v2.0 (Reuscher et al. 2018). Sequences (*O. glaberrima*, *O. glumaepatula*, and *O. sativa* ssp. *japonica*) for each original alignment block were used as a query, and blastn (Altschul et al. 1990) was performed against these genome assemblies. The top hit (*E*-value ≤ 0.00001) was obtained and analyzed.

To assess potential errors in orthologous region assignment of the multiple genome alignment due to hidden paralogous regions, we utilized the high-quality T2T assembly of *O. sativa* ssp. *japonica* cv. Nipponbare (Shang et al. 2023). Using this assembly, the degree of duplication was calculated for each alignment block: the sequence of *O. sativa* ssp. *japonica* in the original alignment block was used as a query, and blastn search was performed against the T2T assembly. The lengths of the blast hits with *E*-value ≤ 0.00001 and with percent identity ≥ 90 were summed and then divided by the length of the query sequence; this ratio was regarded as degree of duplication of each alignment block.

### Overlap with genome annotation

We analyzed the overlap of candidate introgressed regions with annotated genes. Genome annotation data were downloaded from RAP-DB version 2025-03-19 (Sakai et al. 2013). The overlaps of the annotated genes and the genome alignment blocks were identified using the GenomicRanges package (Lawrence et al. 2013) in R version 4.3.3.

Gene ontology (GO) enrichment analysis was performed with agriGO v2.0 (Tian et al. 2017). For each gene set that overlapped with the genome alignment blocks, singular enrichment analysis (SEA) was conducted with default parameters.

### Gene-based analysis

Orthologous gene sets of *Oryza* species were downloaded from Ensembl Plants Genes 62 using BioMart (Kinsella et al. 2011). We used only sequences annotated as one-to-one orthologs. We excluded fragmented sequences shorter than 90% of the corresponding *O. sativa* ssp. *japonica* ortholog in amino acid length. If multiple isoforms exist for a gene, the longest isoform was selected. Amino acid sequences were aligned using the G-INS-i algorithm of MAFFT (Katoh and Standley 2013), and the corresponding nucleotide sequences were aligned based on the amino acid sequences using PAL2NAL (Suyama et al. 2006). A phylogenetic tree for each orthologous gene set was inferred based on the ML method using MEGA version 11.0.13 under the GTR+G+I model with five discrete gamma rate categories and 1000 bootstrap replicates.

### Visualization and statistical test

The results were visualized using the ggplot2 package (Wickham 2016) in R version 4.4.3. The genomic distribution of candidate introgressed regions was visualized using the chromPlot package (Oróstica and Verdugo 2016). Statistical tests were performed with the functions fisher.test, kruskal.test, and wilcox.test in R.

### Generative AI usage

ChatGPT and Gemini were used for English language editing and assistance in coding. All outputs were reviewed and edited by the authors, who take full responsibility for the contents.

## Results

### Phylogenetic incongruence across the genome

To search for genomic regions whose phylogenetic trees showed a topology different from the topology reflecting well-supported consensus species relationships, we first conducted phylogenetic analysis based on a whole-genome alignment of nucleotide sequences from 11 *Oryza* species. Genome alignment blocks which are duplication-free and contain all the species were concatenated, and a phylogenetic tree was inferred based on the ML method, using *O. brachyantha* (FF-genome species) as on outgroup. The resulting topology (Fig. 1a) was consistent with the known relationships among these species, confirming the monophyly of Asian and African (except *O. longistaminata*) species groups and the distant placement of *O. meridionalis* and *O. longistaminata* (Fig 1a).

**Figure 1.**
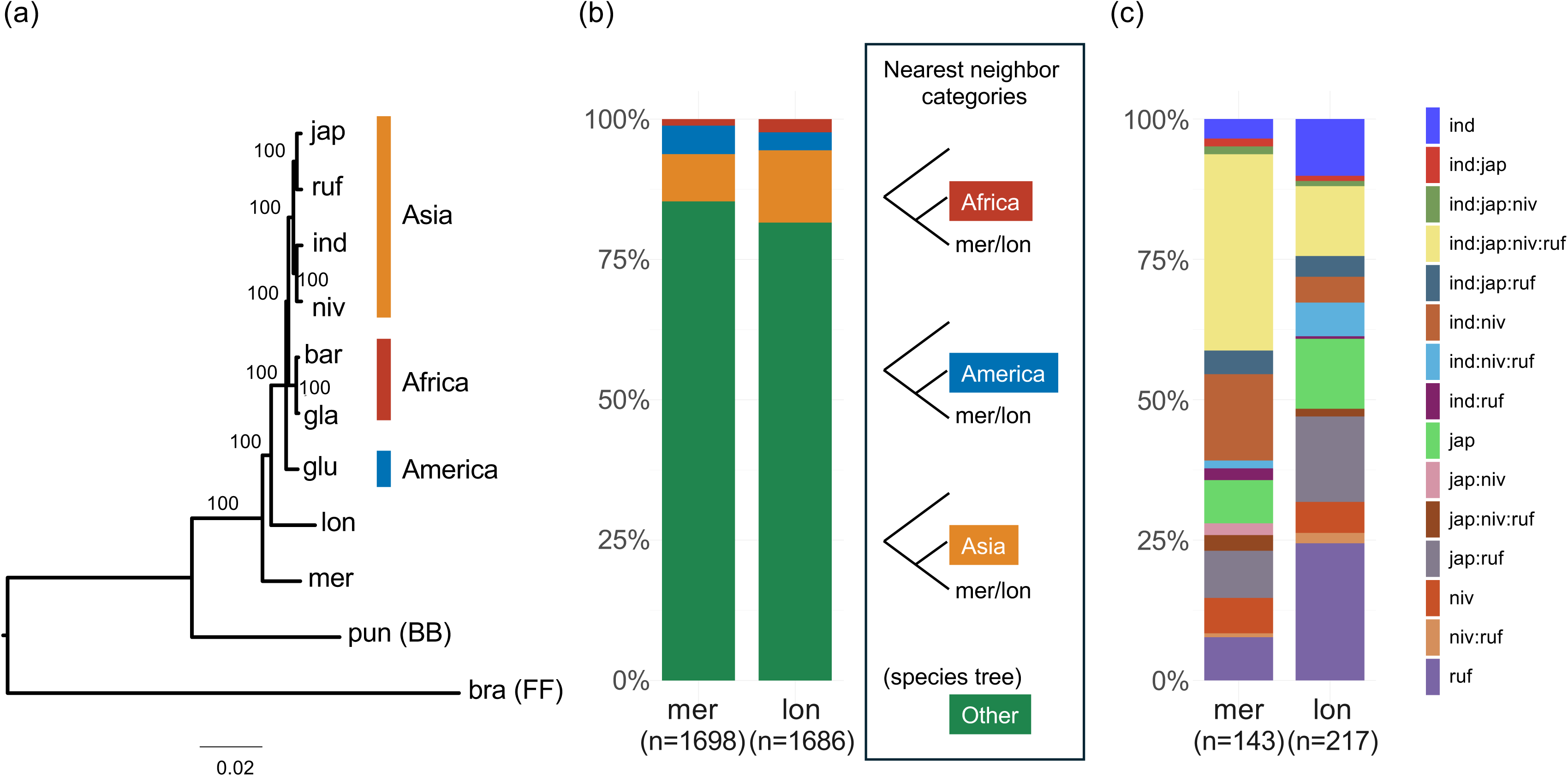
Nearest neighbors of *O. meridionalis* and *O. longistaminata* (a) Phylogenetic tree of *Oryza* species based on a concatenated whole-genome alignment of 17,991,008 sites. Numbers on internal branches indicate bootstrap support (%). (b) Bar chart showing the proportion of individual alignment blocks (trees) classified into four categories based on the nearest neighbors of *O. meridionalis*/*O. longistaminata*. Four categories are Asia (*O. sativa* ssp. *japonica*, *O. rufipogon*, *O. sativa* ssp. *indica*, and *O. nivara*), America (*O. glumaepatula*), Africa (*O. barthii* and *O. glaberrima*) and ‘Other’ (*O. longistaminata*, *O. meridionalis*, and *O. punctata*). For example, ‘Asia’ indicates that the nearest neighbors of *O. meridionalis* or *O. longistaminata* consisted of species only from Asia (*O. sativa* ssp. *japonica*, *O. rufipogon*, *O. sativa* ssp. *indica*, and/or *O. nivara*). When the nearest neighbors included species from more than one categories, it was also classified as ‘Other’. (c) Breakdown of the diversity of Asian species combinations found to be the nearest neighbors within the Asia category in (b). Species abbreviations used throughout this study are as follows: jap: *O. sativa* ssp. *japonica*; ruf: *O. rufipogon*; ind: *O. sativa* ssp. *indica*; niv: *O. nivara*; bar: *O. barthii*; gla: *O. glaberrima*; glu: *O. glumaepatula*; lon: *O. longistaminata*; mer: *O. meridionalis*; pun: *O. punctata*; and bra: *O. brachyantha*.

However, when we examined phylogenetic trees from individual alignment blocks, a different picture emerged (Fig. 1b). Here, we focused on the relationship between most distantly related *Oryza* AA-genome species, *O. meridionalis* and *O. longistaminata*, and classified the alignment blocks (or their trees) based on the nearest neighbors of *O. meridionalis* and *O. longistaminata*. We observed phylogenetic incongruence, concerning the placement of *O. meridionalis* and *O. longistaminata*. Based on the consensus tree topology (Fig. 1a), the nearest neighbors of *O. meridionalis* should be classified as ‘Other’, because the nearest neighbors include species from multiple categories (see below). However, 8.4% of alignment block trees (143 trees) unexpectedly clustered it as the nearest neighbor to the Asian species (*O. sativa*, *O. rufipogon*, and/or *O. nivara*), hereafter referred to as the Asia category. Likewise, 5.1% (86 trees) clustered it with the American species (*O. glumaepatula*; hereafter the America category), while 1.2% (20 trees) clustered it with the African species (*O. barthii* and/or *O. glaberrima*; hereafter the Africa category). When we applied a stringent bootstrap support threshold of ≥ 90 for the cluster, still the Asia category accounted for 5.4% of high-confidence trees (62 trees), while the America and Africa categories accounted for 3.8% (43 trees) and 0.70% (8 trees), respectively. A more pronounced pattern was observed for *O. longistaminata*. The nearest neighbors of *O. longistaminata* should be classified as ‘Other’ for the consensus tree topology (Fig. 1a), but 217 (13%), 54 (3.2%), and 40 (2.4%) trees were classified as Asia, America, and Africa, respectively. When the nearest neighbors with bootstrap support ≥ 90 were selected, we still obtained similar results, in which 152 (15%), 25 (2.4%), and 17 (1.6%) trees were classified as Asia, America, and Africa, respectively. Breakdown of the contents of the Asia category revealed no single species predominated as the nearest neighbors; instead, it comprised a variety of species combinations (Fig. 1c).

### Distinguishing introgression from incomplete lineage sorting

Phylogenetic incongruence could arise from ILS or gene flow such as introgression. To distinguish these possibilities, we performed four-taxon (ABBA-BABA) tests assuming the tree topology (((P1, P2), P3), A). We used *O. punctata* (BB-genome species), the closest relative of the AA-genome species (Fig. 1a), as an outgroup ‘A’. We set either *O. meridionalis* or *O. longistaminata* as P3, and all other possible combinations of the remaining AA-genome species for P1 and P2. Then *D*-statistics were calculated, in which positive values of *D* indicate introgression between P2 and P3, whereas negative values indicate introgression between P1 and P3. The resulting *D*-statistics provided significant evidence for introgression, particularly between *O. meridionalis*/*O. longistaminata* and Asian species (Fig. 2), consistent with the phylogenetic analyses (Fig. 1b). Notably, introgression between *O. longistaminata* and Asian species was prominent, with 17 combinations detected as statistically significant.

**Figure 2.**
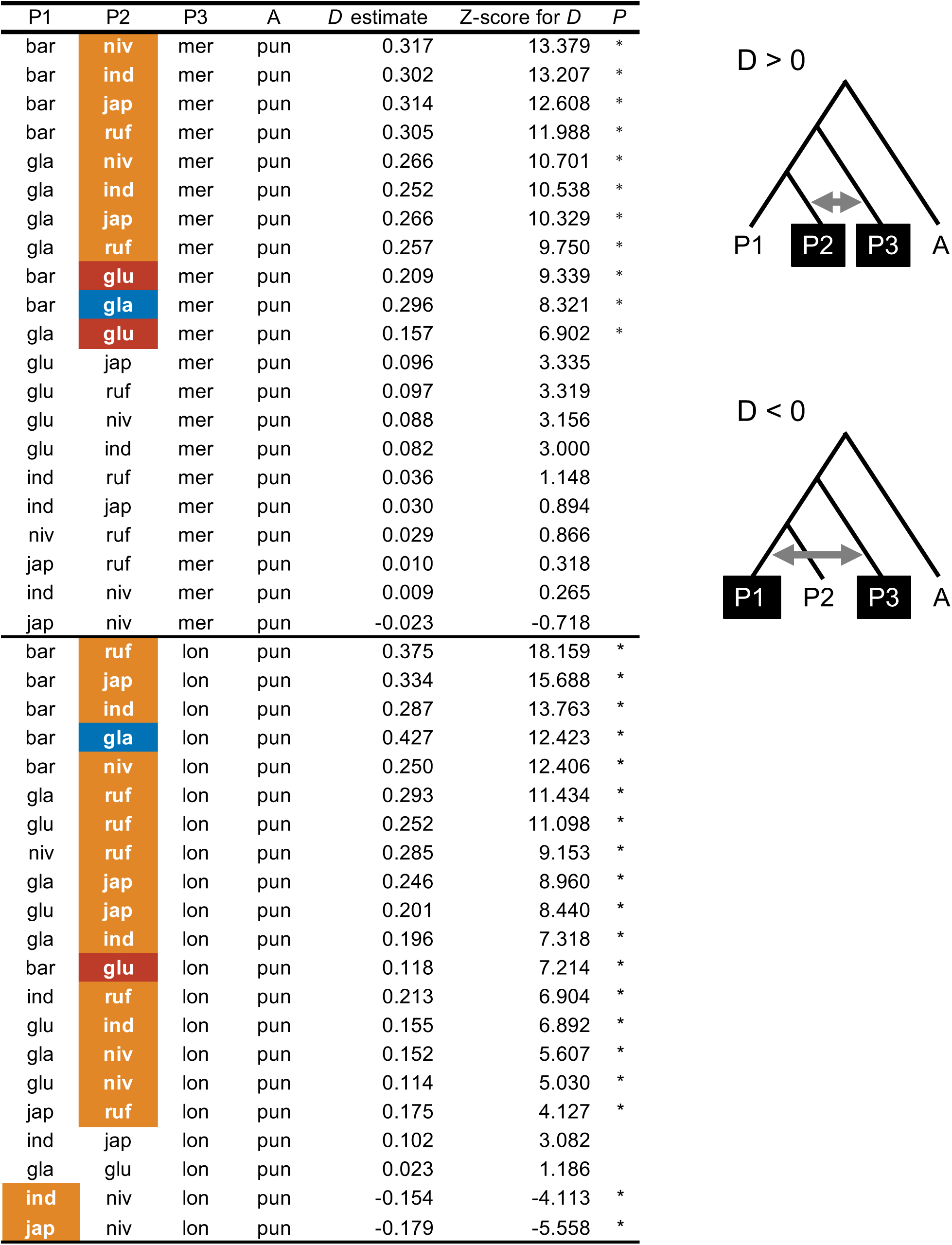
Results of four-taxon (ABBA-BABA) tests. Asterisks indicate *P* ≤ 0.01 after Bonferroni correction for multiple testing. Species showing evidence of introgression with P3 are colored.

### Divergence time estimation

If, for example, the monophyly of *O. longistaminata* and the Asian species resulted from introgression, the divergence time between these species would be expected to be younger than their actual species divergence time. Therefore, the divergence time for each tree of an alignment block was analyzed. We calculated relative times of divergences, without specifying calibration points (Tamura et al. 2012), between *O. meridionalis*/*O. longistaminata* and its nearest neighbors. Using *O. brachyantha* as an outgroup, we calculated the divergence time relative to the deepest node for each tree. The estimated relative divergence times between *O. meridionalis*/*O. longistaminata* and its nearest neighbor species in the Asia/America/Africa category showed younger distribution (*P* ≤ 0.01 by Wilcoxon test, Fig. 3) compared with those in the ‘Other’ category, which included the cases consistent with the consensus tree topology. For example, the mean relative divergence time between *O. longistaminata* and its nearest neighbor species in the Asia category was estimated as 0.134, whereas those of America and Africa categories were estimated as 0.188 and 0.138, respectively. In contrast, the mean relative divergence time between *O. longistaminata* and its nearest neighbor species in the ‘Other’ category was 0.315, showing an older distribution.

**Figure 3.**
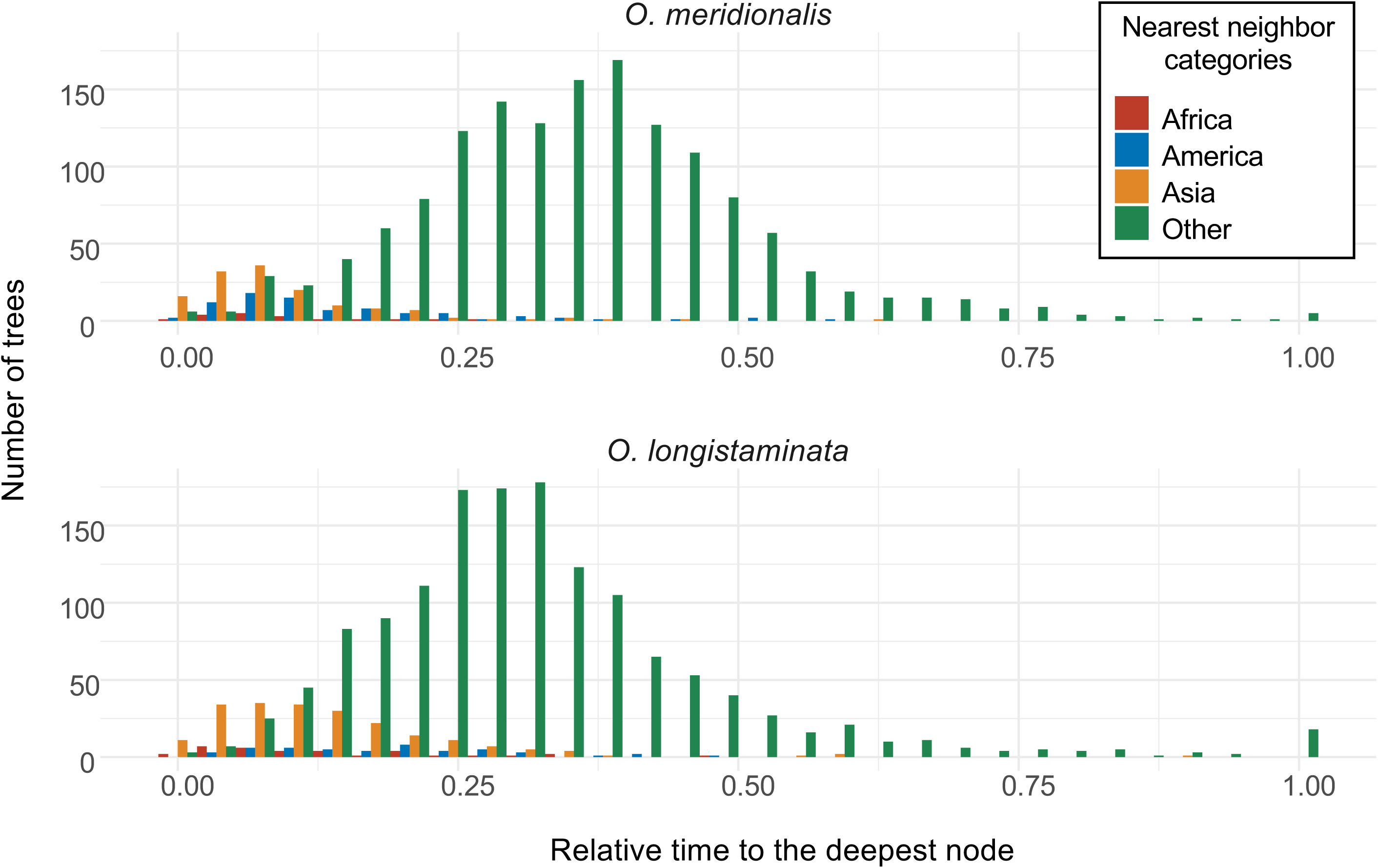
Relative divergence time between *O. meridionalis*/*O. longistaminata* and its nearest neighbors for each category (Africa, America, Asia, and ‘Other’). The estimated relative divergence time for each alignment block was normalized by dividing the relative divergence time of the deepest node in that block’s tree. The resulting values were visualized as histograms.

We also applied a Bayesian approach to estimate the absolute divergence time with the calibration point; 15 million years ago for the divergence of AA/FF genome species (Tang et al. 2010; Stein et al. 2018). First, we estimated the divergence time of the consensus tree topology (Fig. 1a) based on the whole-genome alignment. The species divergence time between *O. meridionalis* and other species was 1.78 million years ago (95% HPD: 1.22–2.39) while that of *O. longistaminata* and other species was estimated as 1.40 million years ago (95% HPD: 0.933–1.89) (Supplementary Fig. 1). In contrast, the median divergence time for the incongruent alignment blocks of the Asia category was 0.319 million years ago (0.441 ± 0.359) for *O. meridionalis* and 0.536 million years ago (0.584 ± 0.345) for *O. longistaminata*, younger than the speciation times estimated based on the whole-genome alignment. Since older divergence time is expected for ILS, these results therefore suggested that a part of phylogenetic incongruence could be explained by gene flow such as introgression. We hereafter refer to these alignment blocks assigned to Asia, America, and Africa nearest neighbor categories as candidate introgressed regions.

### Assessment of sequence features for potential technical artifacts

Phylogenetic incongruence can be explained not only by biological factors but also by analytical factors (Fleming et al. 2023; Steenwyk et al. 2023). To search for possible analytical factors for the observed phylogenetic incongruence, we examined several sequence features and compared them among the nearest neighbor categories (Africa, America, Asia, and Other).

First, we tested homogeneity for nucleotide composition for all sequences in each alignment block with IQ-TREE. As a result, composition heterogeneity (*p* < 0.05 in the IQ-TREE chi-squared test) was found fewer than 10% of the alignment blocks (Fig. 4a). Species showing composition heterogeneity were *O. brachyantha* (81%), *O. punctata* (11%), and *O. longistaminata* (6.5%), possibly reflecting the distant relationships with AA-genome species. We however found no enrichment of heterogeneity (*P* ≥ 0.01 by Fisher’s exact test) for the candidate introgressed regions (Africa, America, and Asia categories).

**Figure 4.**
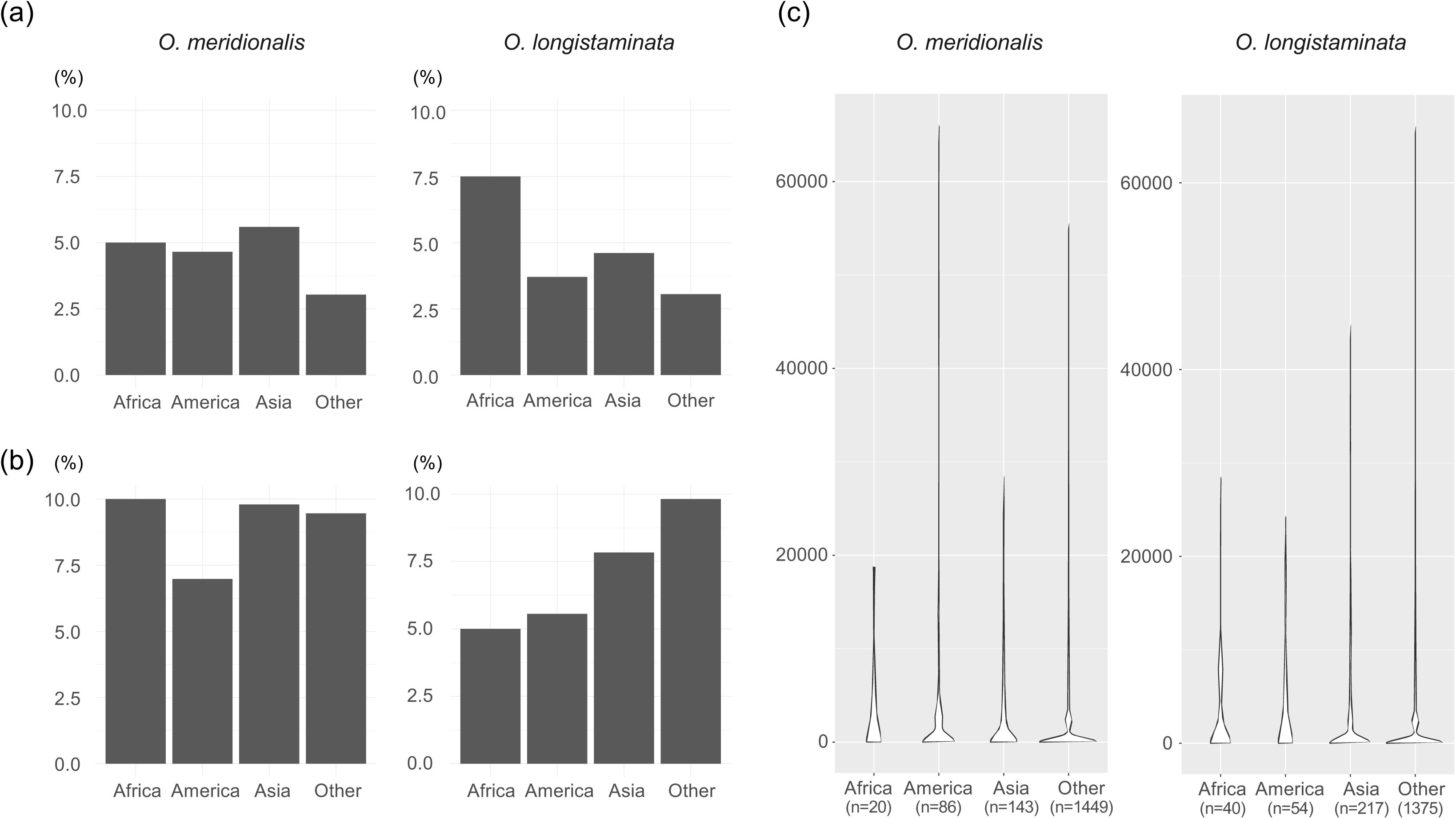
Assessment of sequence features for potential technical errors grouped by the nearest neighbor categories (Africa, America, Asia, and ‘Other’) of *O. meridionalis*/*O. longistaminata*. (a) Bar chart showing the proportion of alignment blocks exhibiting significant compositional bias among sequences. (b) Bar chart showing the proportion of alignment blocks for which the assumptions of stationarity and/or hovmogeneity were rejected. (c) Violin plot showing the distribution of C values 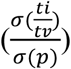, defined as the ratio of the standard deviation of transition-to-transversion ratios (*ti*/*tv*) to the standard deviation of the genetic distance (*p*), which serves as an indicator of nucleotide substitution saturation (Struck et al. 2008).

Second, to assess whether the sequence data violated the assumption of substitution symmetry, a test of symmetry was performed with IQ-TREE. The proportion of the alignment blocks for which the assumptions of stationarity and/or homogeneity were rejected (*P* ≤ 0.01) was below 10% (Fig. 4b). We again found no enrichment of violation (*P* ≥ 0.01 by Fisher’s exact test) for the candidate introgressed regions.

Third, site saturation of nucleotide substitution, especially for outgroup species, may affect phylogenetic inferences. Therefore, we examined C value, in which small values indicate saturation (Struck et al. 2008). As a result, the median C values were 11.3, 10.0, 12.9, and 9.87 for the Africa, America, Asia, and ‘Other’ categories of *O. meridionalis*, respectively (Fig. 4c, left). For *O. longistaminata*, the corresponding values were 10.1, 15.1, 8.57, and 10.2, respectively (Fig. 4c, right). We found no statistically significant differences among the categories (*P* ≥ 0.01 by Kruskal-Wallis test).

### Validation with other genome assemblies

Another possible analytical error could be the primary dataset of genome alignment used. To rule out issues with the dataset used, we assessed our findings using alternative genome assemblies available for *O. meridionalis* (Abdullah et al. 2025) and *O. longistaminata* (Reuscher et al. 2018; Lian et al. 2024; Guang et al. 2025). We compared the sequences of these assemblies with those used in the primary dataset: using sequences (*O. glaberrima*, *O. glumaepatula*, and *O. sativa* ssp. *japonica*) of each alignment block in the original dataset as a query, we performed sequence similarity search against the alternative genome assemblies and obtained sequences with the highest similarity. These analyses confirmed that the candidate introgressed regions showed higher sequence similarity than the alignment blocks in ‘Other’ category (*P* ≤ 0.01 by Wilcoxon test, Fig. 5). For example, the mean sequence identities between *O. sativa* ssp. *japonica* in the alignment blocks of Asia category and three alternative assemblies of *O. longistaminata* were 97.153–97.396, which were higher than 96.368-96.471 in those of ‘Other’ category (P ≤ 0.01 by Wilcoxon test). A similar pattern was observed for queries of *O. glumaepatula* in the America category and queries of *O. glaberrima* in the Africa category. All these results were consistent with possible introgression between these species regardless of genome assemblies used.

**Figure 5.**
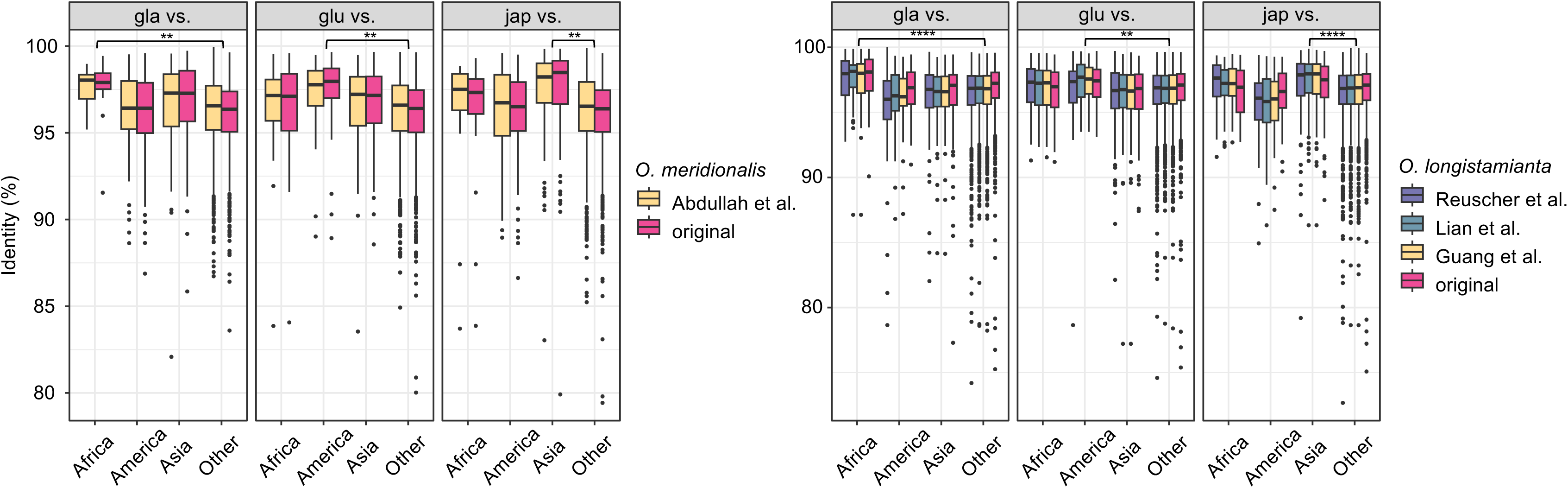
Box plot showing sequence identity between the sequences in the alignment blocks (grouped by the nearest neighbor categories) and the sequences from alternative genome assemblies of *O. meridionalis*/*O. longistaminata*. For each alignment block, the sequences of *O. glaberrima*, *O. glumaepatula*, and *O. sativa* ssp. *japonica* were used as queries, serving representatives of the Africa, America, and Asia categories, respectively. ‘Original’ indicates the sequences of *O. meridionalis*/*O. longistaminata* from the alignment blocks (the primary dataset). The numbers of dataset for each category are the same as in Fig. 4c. Asterisks indicate the number of pairs that are significantly different (*P* ≤ 0.01 by Wilcoxon test) between the following category pairs: Africa and ‘Other’ for the *O. glaberrima* queries, America and ‘Other’ for the *O. glumaepatula* queries, and Asia and ‘Other’ for the *O. sativa* ssp. *japonica* queries. Boxes represent the interquartile range (IQR), the center line indicate the median, whiskers extend to 1.5 × IQR, and outliers are shown as individual points.

Finally, we evaluated potential mis-assignment of orthologous regions in the original multiple alignment, due to hidden paralogous regions. Although the reference assembly used in the multiple alignment was IRGSP-1.0 of *O. sativa* ssp. *japonica* cv. Nipponbare, a new high-quality T2T assembly of *O. sativa* ssp. *japonica* cv. Nipponbare was published (Shang et al. 2023). Thus, we searched the *O. sativa* ssp. *japonica* sequence of each alignment block for the new T2T assembly, and we calculated the degree of duplication including repetitive sequences, which may affect genome assembly and alignment. As a result, the median degree of duplication was 2.41, 5.78, 4.45, and 7.05 for the Africa, America, Asia, and ‘Other’ categories of *O. meridionalis*, respectively (Fig. 6). We found no statistically significant differences among the categories (*P* ≥ 0.01 by Kruskal-Wallis test). For *O. longistaminata*, the corresponding values were 2.99, 3.28, 5.98, and 7.21 respectively. The difference was significant at 0.001. However, when we selected only the nearest neighbors with bootstrap support ≥ 90, the difference was not significant (*P* ≥ 0.01 by Kruskal-Wallis test).

**Figure 6.**
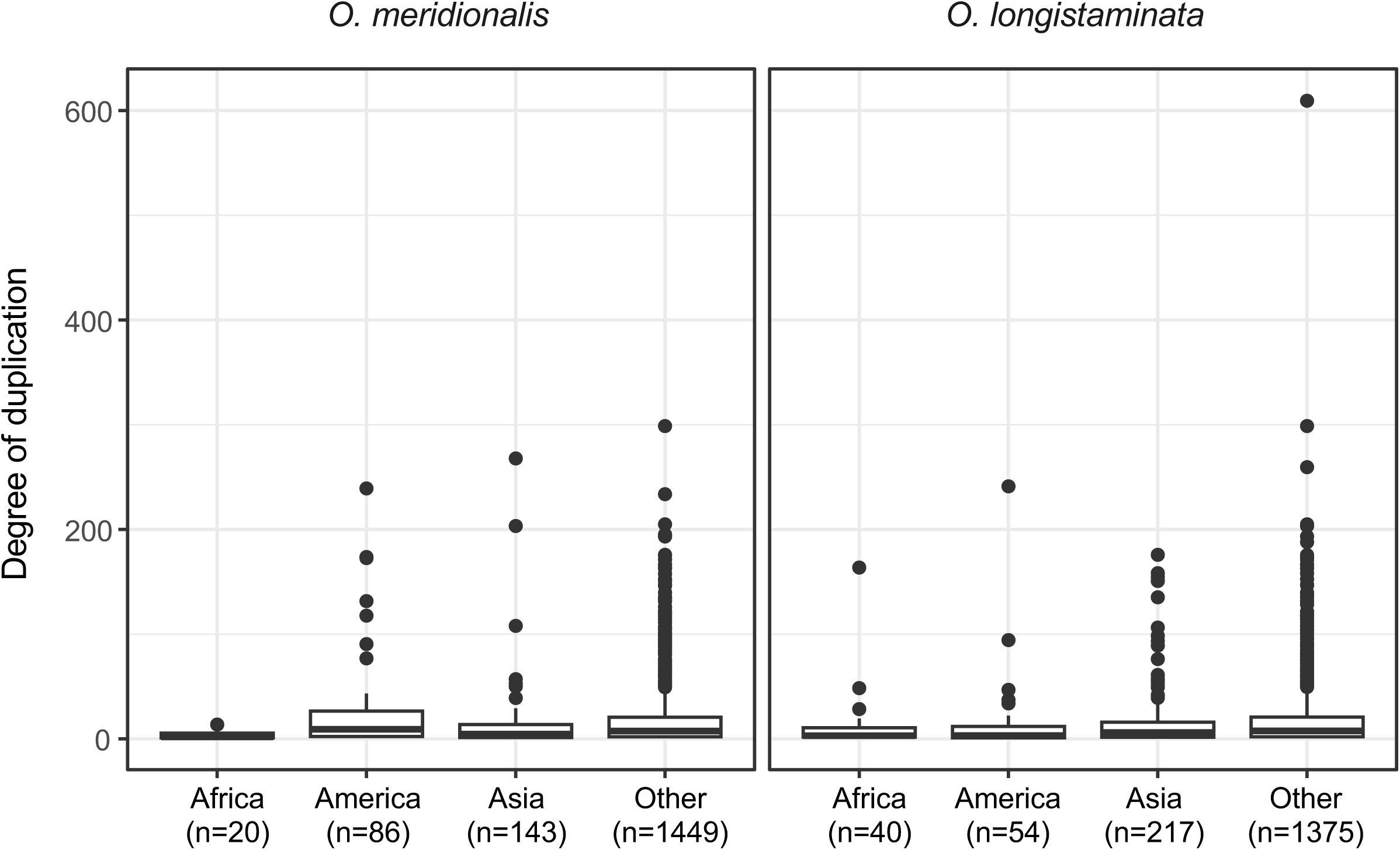
Degree of duplication for the alignment blocks grouped by the nearest neighbor categories (Africa, America, Asia, and ‘Other’) of *O. meridionalis*/*O. longistaminata*. The degree of duplication was calculated as the total length of homologous segments in the T2T assembly of *O. sativa* ssp. *japonica* (Shang et al. 2023) divided by the length of the query sequence (*O. sativa* ssp. *japonica* IRGSP v1.0) for each alignment block. Boxes represent the interquartile range (IQR), the center line indicate the median, whiskers extend to 1.5 × IQR, and outliers are shown as individual points.

### Characteristics of the candidate introgressed regions

The candidate introgressed regions appeared to be randomly distributed across the chromosomes, with no obvious clustering (Supplementary Fig. 2). We also examined overlaps with genic regions based on *O. sativa* ssp. *japonica* IRGSP 1.0 genome annotation. Within the high-confidence (bootstrap support ≥ 90) candidate introgressed regions, 149, 87, and 5 genes were overlapped with the genome alignment blocks of Asia, America, and Africa categories of *O. meridionalis*, while 291, 75, and 48 genes were overlapped with the genome alignment blocks of Asia, Africa, and America categories of *O. longistaminata*. To assess potential functional enrichment, we performed GO enrichment analysis of the genes within these candidate introgressed regions. We found no statistically significant functional bias. Phylogenetic analysis of orthologs of these candidate introgressed genes showed various tree topologies. We found two apparent examples showing a cluster of *O. longistaminata* and Asian species (Fig. 7).

**Figure 7.**
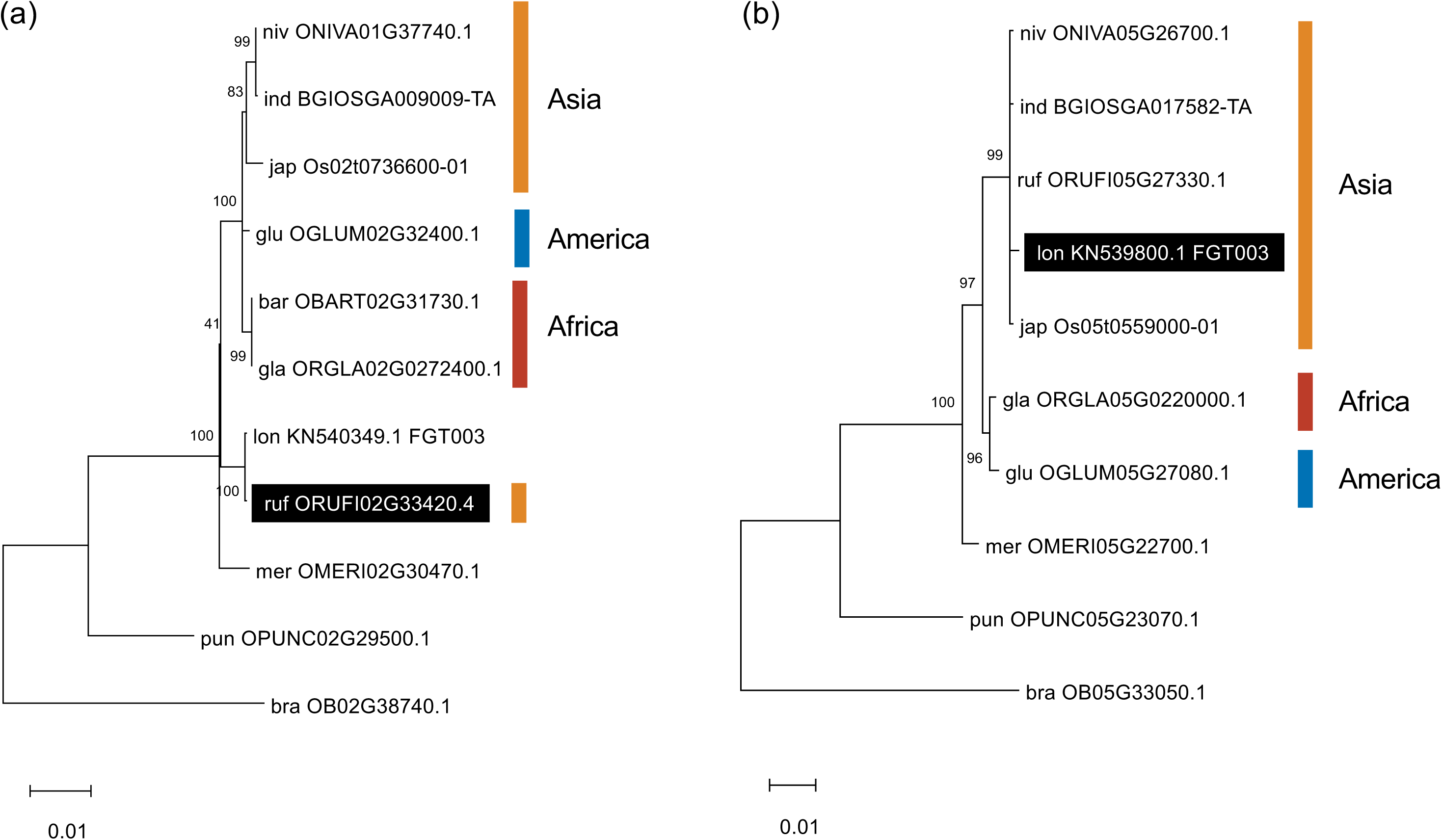
Examples of phylogenetic trees for candidate introgressed genes. The trees for (a) DEAD-like helicase, N-terminal domain containing protein and (b) Galactose oxidase/kelch, beta-propeller domain containing protein were inferred using the ML method, based on 3381 and 1608 nucleotide sites, respectively. Numbers on internal branches indicate bootstrap support (%). The candidates for introgressed sequences are indicated in reverse text.

## Discussion

In this study, we identified phylogenetic incongruence between alignment blocks of distantly related *Oryza* AA-genome species, especially *O. meridionalis*/*O. longistaminata* and other species (Fig. 1). Divergence times of these incongruent alignment blocks showed a younger divergence time compared with those of other alignment blocks (Fig. 3), suggesting introgression between these species. Four-taxon ABBA-BABA test also supported this could be explained by introgression but not by ILS (Fig. 2). Phylogenetic incongruence can arise not only from biological factors but also from analytical factors (Fleming et al. 2023; Steenwyk et al. 2023). Therefore, we examined the sequence features of these candidate introgressed regions. We found no evidence of potential technical artifacts, at least with respect to compositional bias (Fig. 4a), symmetry assumptions (Fig. 4b), sequence substitution saturation (Fig. 4c), and the degree of duplication (Fig. 6), although a lack of statistical significance should not be interpreted as evidence for the null hypothesis. We cannot fully exclude the possibility that analytical factors may have influenced our results, nevertheless, these lines of evidence suggest the existence of genomic footprints of historical introgression between ancient lineages of wild *Oryza* AA-genome species that are now separated by vast geographical distances. Our finding of introgression between *O. meridionalis* and Asian species (especially *O. rufipogon* whose geographical distributions overlap with that of *O. meridionalis* in Australia) is consistent with previous reports of historical and ongoing gene flow between them (Brozynska et al. 2017; Moner, Furtado, Chivers, et al. 2018; Fujino et al. 2019; Hasan et al. 2022). This consistency validates our study.

The novel finding in this study provides genome-wide evidence for historical introgression between ancient lineages of African species *O. longistaminata* and Asian species. This is particularly surprising given their current distinct geographical distributions. *O. longistaminata* is a perennial and self-incompatibility species, and hybridization with *O. sativa* is thought to be limited (Tong et al. 2023; Mangosongo et al. 2024), for which molecular mechanisms of hybrid sterility have been studied (Tonosaki et al. 2018; Myint et al. 2024). However, our finding aligns with a recent report of natural hybridization and bidirectional introgression between *O. sativa* and *O. longistaminata* in Southern Africa based on RAD-seq data (Labroo et al. 2023). Furthermore, incongruent phylogenies between nuclear and organellar genomes have long been reported for AA-genome species (Duan et al. 2007; Kim et al. 2015; Yin et al. 2015; Fornasiero et al. 2025). Chloroplast is known to be easily affected by gene flow, known as chloroplast capture (Rieseberg 1991; Tsitrone et al. 2003), and reticulate evolution prior to rice domestication was suggested (Moner, Furtado, and Henry 2018; Moner et al. 2020). Therefore, this study uncovered genomic footprints within the nuclear genome, providing direct evidence of ancient, reticulate evolutionary events.

Detection of various combinations of Asian species clustering with *O. longistaminata* (Fig. 1c) suggests a scenario involving multiple, and possibly bidirectional, introgression events between ancestral lineages of these species. In fact, introgression from *O. longistaminata* to Asian species (Fig. 7a) and also from Asian species to *O. longistaminata* (Fig. 7b) were inferred, suggesting bidirectional introgression. The median divergence time of candidate introgressed regions between *O. longistaminata* and the Asian species was estimated as 0.536 million years ago. In comparison, the earliest divergence among Asian rice lineages was estimated as 0.35 million years ago in this study (Supplementary Fig. 1) and as 0.55–0.99 million years ago in previous studies (Zhu et al. 2014; Brozynska et al. 2017; Stein et al. 2018). Therefore, introgression between these species might be around the time of the speciation of the Asian species. It remains unclear whether reproductive barriers were fully established in the past, and hybridization may therefore have been possible during this period. It is noteworthy that the estimated time does not necessarily reflect the actual timing of introgression, because the species used in this study may not be a direct progenitor of introgression. For example, it is possible that an ancient species related to current *O. longistaminata*, which already was extinct or remains unsampled, might have hybridized with ancient lineages of Asian species, a process known as ghost introgression (Ottenburghs 2020).

Identifying the geographical location of introgression events between ancient lineages of African *O. longistaminata* and the Asian species is challenging. The intercontinental divergence of AA-genome species occurred long after the breakup of Gondwanaland, and long-distance dispersal mediated by animals has been hypothesized, either from an Asian origin (Vaughan et al. 2005; Tang et al. 2010; Lin et al. 2015) or from an African origin (Cheng et al. 2002). Since the geographical distribution of ancient AA-genome species at the time of introgression is unknown, two possibilities can be considered: these species may already have been geographically separated (allopatric) as observed today, or they may have overlapped geographically (sympatric). The former possibility seems unlikely given the limited dispersal ability of pollen of rice (Song et al. 2003). Under the latter possibility, introgression may have occurred with a now-extinct ancestral lineage of *O. longistaminata* in Asia (i.e., ghost introgression). Interestingly, the possible existence of an Asian species related to *O. longistaminata* has been reported (Htut et al. 2021; Guo et al. 2025), which may be consistent with our finding. Trans-continental hybridization has been documented in the genus *Gossypium* (Wendel and Cronn 2003; Xu et al. 2024) and also between American *O. glumaepatula* and African *O. barthii* (Stein et al. 2018). These suggest that gene flow across continents might have played an important role in the evolutionary history of *Oryza*.

This study has some limitations, which suggest directions for future research. First, our analysis depends on a specific set of publicly available genome assemblies and their alignment. We limited the genomic regions where sequences of all the 11 species were available, which could have underestimated the amount of candidate introgressed regions. In addition, our analyses were based on alignment blocks, reflecting synteny blocks across species (Herrero et al. 2016). Precise identification of an introgressed segment within each alignment block could refine estimates of divergence times. Second, the use of a single representative genome per species does not account for the considerable genetic diversity that exists within species populations. Although whole-genome genetic variation data of many accessions are available, especially for cultivars (Huang et al. 2012; Ohyanagi et al. 2016; Cubry et al. 2018; Wang et al. 2018; Guo et al. 2025), those derived from natural populations of *O. meridionalis* and *O. longistaminata* are still limited, despite the high diversity reported for these species (Kiambi et al. 2005; Moner, Furtado, Chivers, et al. 2018; Melaku et al. 2019; Getachew et al. 2020; Lakew et al. 2021; Hasan et al. 2022). Future research incorporating population-level diversity data and employing sophisticated methods like multispecies coalescent models (Xu and Yang 2016; Tiley et al. 2020) will be essential for building a more precise picture of the complex evolutionary history of AA-genome species. Finally, although we did not detect any functional enrichment in the candidate introgressed genes, further investigation of these genes may be warranted.

## Data Availability

All data used in this study are derived from publicly available sources, as described in the main text.

## Acknowledgements

We are grateful to T. Itoh and M. Kumagai for valuable discussion and comments.

## Author Contributions

Y. Kishima conceived the study. K.O.K. designed the study. K.O.K. and Y. Kotoku performed the analyses. K.O.K. and Y. Kishima wrote the manuscript. All authors read, revised, and approved the final manuscript.

