## Supplementary Figures 1 and 2 for "Genomic footprints of historical introgression between ancient lineages of wild *Oryza* AA-genome species with widely separated contemporary distributions"

**Supplementary Materials**

**Supplementary Figure 1.** Estimated divergence time of AA-genome species. Numbers on the nodes represent time in million years ago (MYA). Blue bars indicate the 95% highest posterior density (HPD) intervals.

**Supplementary Figure 2.** Genomic distribution of candidate introgressed regions. The plots show the genomic location of alignment blocks (bootstrap support $\geq$ 90) where the nearest neighbors of (a) *O. meridionalis* and (b) *O. longistaminata* are classified into the four categories (Africa, America, Asia, and ‘Other’).

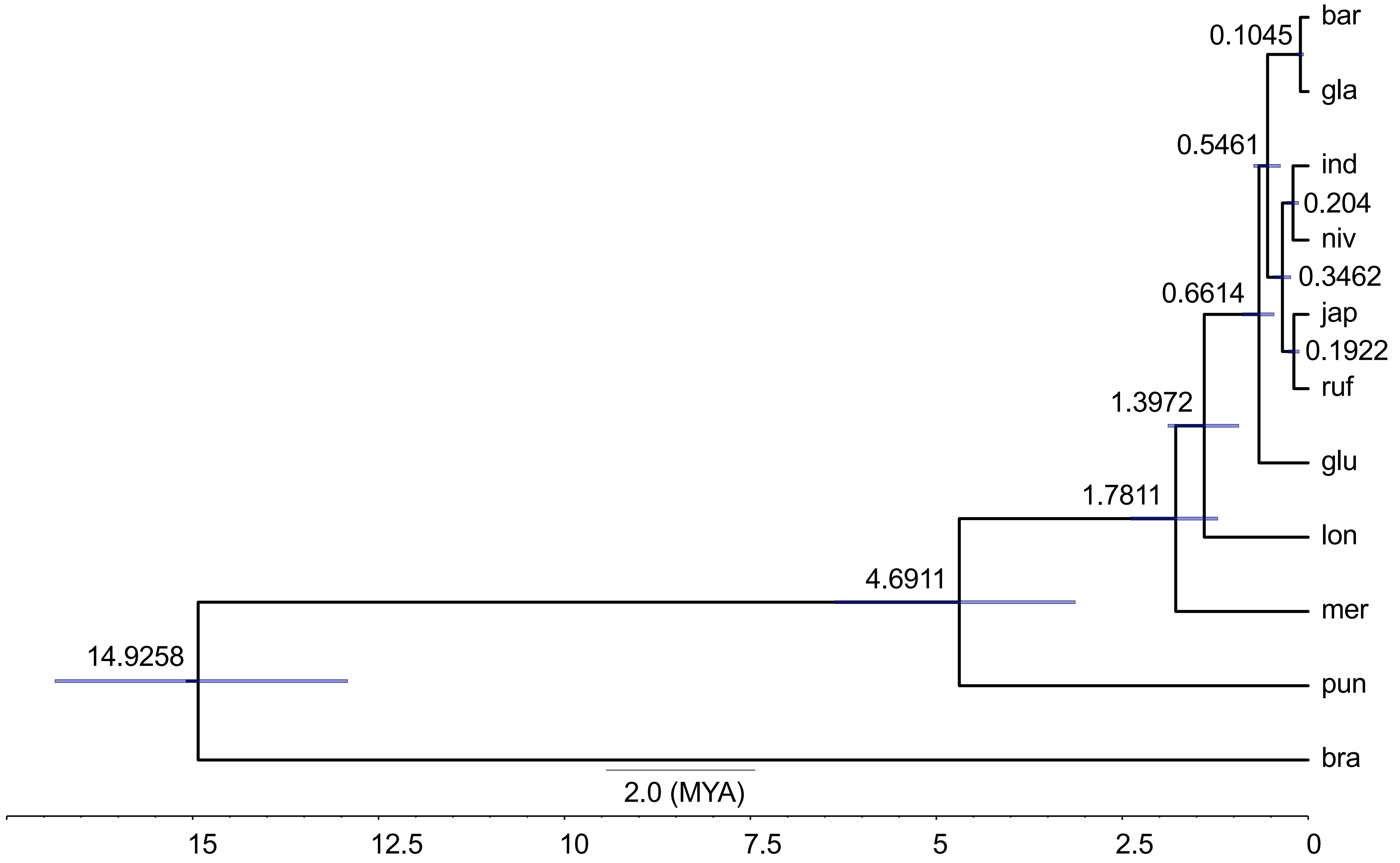


Supplementary Fig. 1

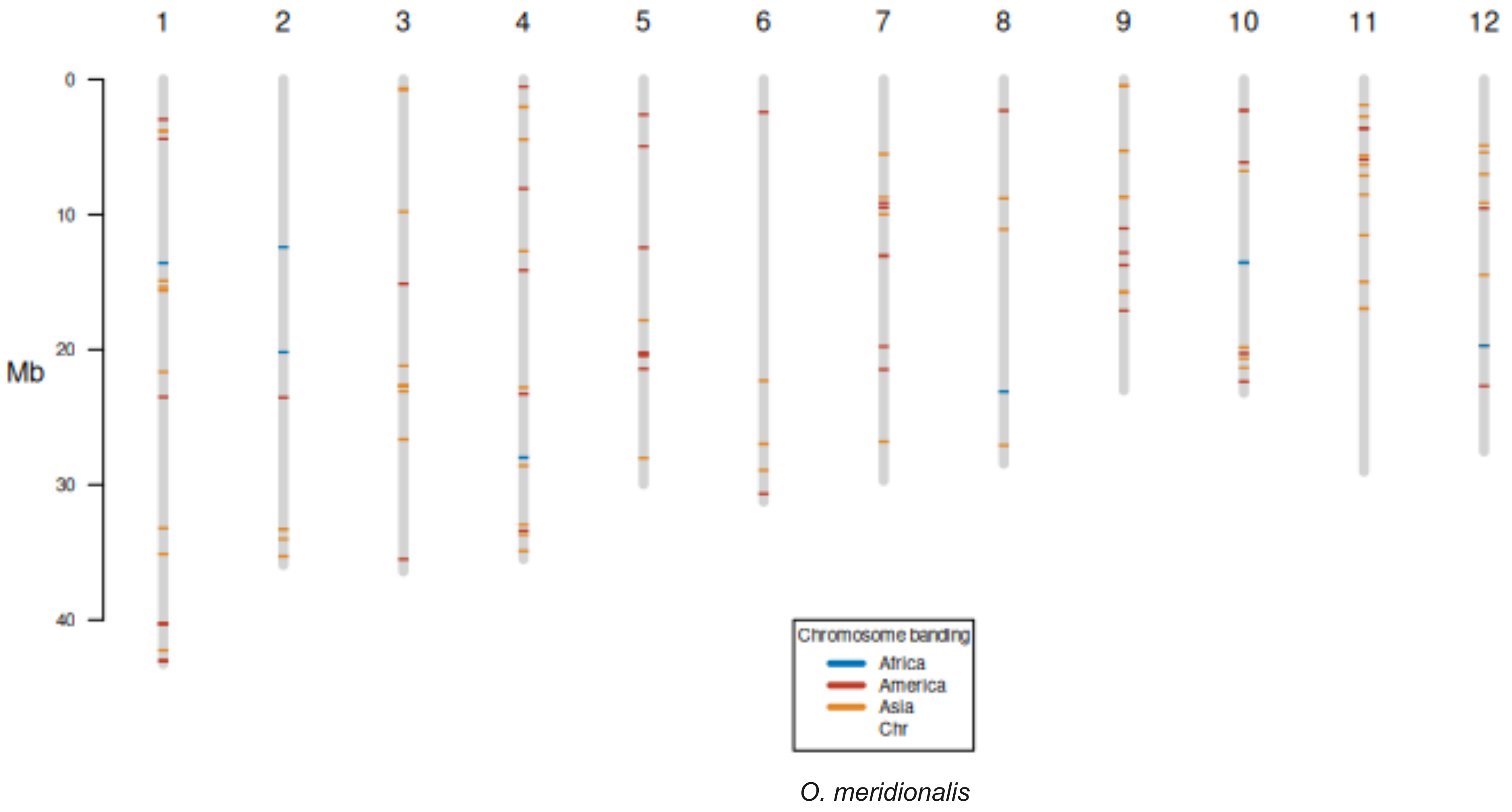


Supplementary Fig. 2a

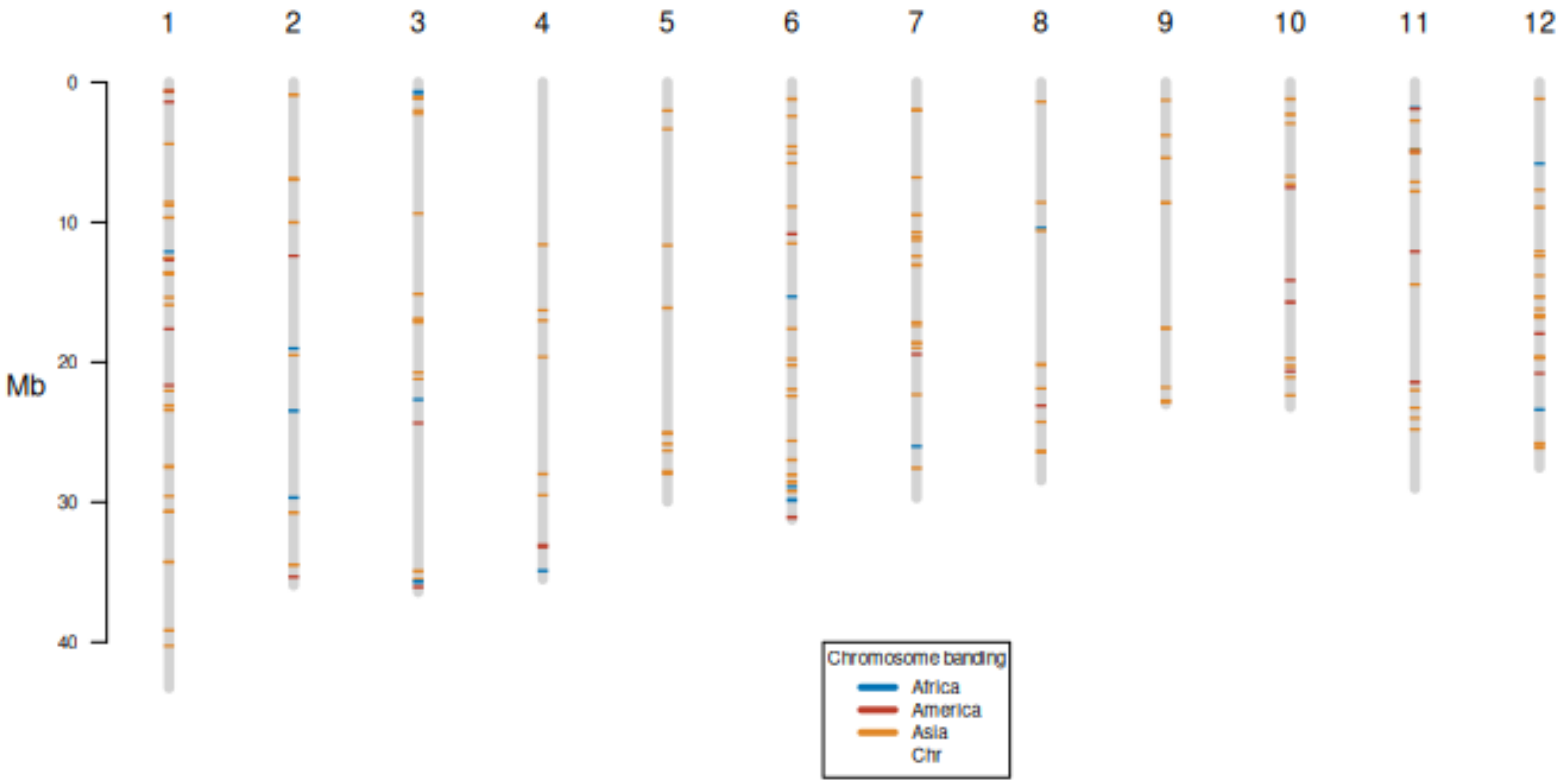


*O. longistaminata*

Supplementary Fig. 2b
